# The Epidermal Barrier is Indispensable for Systemic Energy Homeostasis

**DOI:** 10.1101/2020.09.01.277723

**Authors:** Vibeke Kruse, Ditte Neess, Ann-Britt Marcher, Mie Rye Wæde, Julie Vistisen, Pauline M. Møller, Rikke Petersen, Jonathan R. Brewer, Tao Ma, Georgia Colleluori, Ilenia Severi, Saverio Cinti, Zach Gerhart-Hines, Susanne Mandrup, Nils J. Færgeman

**Affiliations:** Department of Biochemistry and Molecular Biology, Villum Center for Bioanalytical Sciences, University of Southern Denmark, Campusvej 55, 5230 Odense M, Denmark; Novo Nordisk Foundation Center for Basic Metabolic Research, University of Copenhagen, 2200 Copenhagen, Denmark; Center for the study of Obesity, Polytechnic University of Marche, 60020 Ancona, Italy

**Keywords:** Epidermal barrier, energy expenditure, adipose tissue, browning, diet induced obesity, β-adrenergic signaling

## Abstract

**Objectives:** Homeostatic regulation of body temperature is fundamental to mammalian physiology and is controlled by acute and chronic responses of local, endocrine and neuronal regulators. Although the skin is the largest sensory organ of the human body, and plays a fundamental role in regulating body temperature, it is surprising that adaptive alterations in skin functions and morphology only vaguely have been associated with physiological responses to cold stress or sensation of ambient temperatures.

**Methods:** To unravel the physiological responses to a compromised epidermal barrier in detail we have used animal models with either defects in skin lipid metabolism (ACBP^-/-^ and skin-specific ACBP^-/-^ knockout mice) or defects in skin structural proteins (*ma/ma Flg*^*ft/ft*^). The primary objective was to clarify how defects in epidermal barrier function affect 1) energy expenditure by indirect calorimetry, 2) response to high fat feeding and a high oral glucose load and 3) expression of brown-selective gene programs by quantitative PCR in inguinal WAT (iWAT).

**Results:** We show that mice with a compromised epidermal barrier function exhibit increased energy expenditure, increased food intake, browning of the iWAT, and resistance to diet-induced obesity. The metabolic phenotype, including browning of the iWAT, is reversed by housing the mice at thermoneutrality (30°C) or by pharmacological β-adrenergic blocking. These findings show that a compromised epidermal barrier induces a β-adrenergic response that increases energy expenditure and browning of the white adipose tissue to maintain a normal body temperature.

**Conclusion:** Our findings show that the epidermal barrier plays a key role in maintaining systemic metabolic homeostasis.

**Highlights:** Energy expenditure is significantly augmented in mice with impaired epidermal barrier.

Mice with compromised barrier display increased food intake while maintaining normal bodyweight.

Mice with an impaired epidermal barrier are resistant to diet-induced obesity and insulin resistance.

Compromised barrier function induces expression of brown-selective gene programs in iWAT.

Thermoneutral housing or blocking β-adrenergic signaling prevents induction of brite-selective genes in iWAT and reverses food intake.

## 1. INTRODUCTION

The skin is the largest sensory organ of the human body, and accounts for approximately 16% of total body weight. The skin barrier not only protects us from environmental xenobiotics, oxidative stress and infections from various pathogens, it also regulates the amount of heat and water released from the body, and senses various environmental alterations including thermal variations [1]. Mammalian skin is comprised of three layers, the hypodermis, the dermis and the epidermis. The epidermis is further subdivided into multiple layers, among which the outer most layer, the stratum corneum, provides the epidermal permeability barrier. It consists of terminally differentiated keratinocytes, which are embedded in a highly ordered extracellular lipid-containing matrix. Numerous factors, including structural and junctional proteins, serve fundamental roles both in generation and maintenance of the epidermal permeability barrier including terminal epithelial differentiation products such as filaggrin, loricrin, certain keratins and tight junctions. Likewise, lipids like ceramides, sterols and fatty acids also make up crucial constituents of the epidermal barrier [2; 3]. Accordingly, lipid metabolism serves fundamental roles in generation and maintenance of the epidermal permeability barrier and proper insulation [2]. Hence, functional loss of enzymes and transport proteins involved in lipid metabolism can result in severe desiccation and consequently neonatal or premature death. However, non-lethal phenotypes including tousled and greasy fur, increased trans-epidermal water loss (TEWL) and alopecia have also been observed, e.g. in mice lacking stearoyl-CoA desaturase 1 (SCD1) [4], very long chain fatty acid elongase 3 (ELOVL3^-/-^) [5; 6], or the alkaline ceramidase ACER1 [7]. Similarly, we have previously shown that acyl-CoA binding protein (ACBP) not only serves key functions in lipid metabolism in mammals [8; 9] but also that ACBP in keratinocytes is required to maintain an intact epidermal barrier and normal fur and to prevent alopecia [10]. Furthermore, skin-specific loss of ACBP also increases lipolysis in white adipose tissue (WAT) and changes hepatic lipid metabolism [11], indicating that lipid metabolism in the skin affects systemic metabolism. Intriguingly, the effects on lipid metabolism in adipose and liver tissues can be rescued by applying an artificial barrier to the skin of ACBP^-/-^ mice [11], indicating that it is the epidermal barrier function *per se* rather than lipid-derived signaling molecules that lead to the system-wide metabolic effects.

These findings prompted us to investigate how impairment of the epidermal barrier function affects whole body energy metabolism. We show that mice with constitutive knockout of ACBP (ACBP^-/-^) as well as mice with keratinocyte specific knockout of ACBP mice (K14-ACBP^-/-^) exhibit increased energy expenditure, browning of white adipose tissue and resistance to high fat diet-induced obesity. Importantly, we show that these systemic effects on energy metabolism are recapitulated in the flaky tail mice (*ma/ma Flg*^*ft/ft*^), a widely used as a model of human atopic dermatitis. Collectively, our findings show that even modest changes in the epidermal barrier can have major effects on systemic energy homeostasis.

## 2. METHODS

### 2.1 Animal experiments

Mice with constitutive- and conditional targeting of the *Acbp* gene have been described previously [11; 12]. The *ma/ma Flg*^*ft/ft*^ mice were obtained from The Jackson Laboratories and were back crossed to wildtype C57BL/6J mice for at least ten generations to obtain a congenic background. Unless otherwise stated, mice were group housed on a 12-hour light/dark cycle and given ad libitum access to water and chow food (Altromin 1324). For HFD feeding D12492 (Research Diets) was administered ad libitum. Housing at either thermoneutral temp (30°C) or cold exposure (4°C) were performed with individually housed mice for 72 hours. Propranolol injections (20 mg/kg) were administered by I.P. injection once every day for 5 days in mice housed at 22°C. All animal experiments and breeding of transgenic mice were approved by the Danish Animal Experiment Inspectorate.

### 2.2 Real-Time PCR

Frozen iWAT was homogenized in TRI reagent (Sigma) and RNA was isolated according to manufacturer’s protocol. RNA concentration and quality were evaluated by use of NanoDrop™ 1000 spectrophotometer and by gel electrophoresis. RNA was treated with DNase I (Invitrogen), and cDNA was prepared as described previously [12]. mRNA expression levels were determined by real-time PCR and normalized to *Tfiib* expression. All quantitative PCR analyses were carried out on LightCycler^®^ 480 Multiwell Plates 384. Primers used are listed in table S1.

### 2.3 Western blot analysis

Protein extracts were prepared and analyzed as previously described [12]. Primary antibodies used were anti-UCP1 (1:1000) (ab10983, Abcam) and anti-actin (1:1000) (A5441, Sigma). Actin was used as a loading control. The secondary antibody was horseradish peroxidase-conjugated goat anti rabbit IgG (1:2000) (Dako).

### 2.4 Cold exposure

Mice were individually caged and housed at 4°C for 3 days with *ad libitum* access to chow food and water. The mice were sacrificed by anesthesia (25% hypnorm, 25% dormicum in sterile water, 0.01 m/g mouse) followed by blood collection by heart puncture and transcardiacal perfusion with PBS or 4% PFA depending on following experiments.

### 2.5 Sample preparation for microscopy

Mice were anesthetized and transcardially perfused with PBS followed by 4% PFA in 0.1M phosphate butter, pH 7.4. The iWAT was dissected and fixed over night at 4°C. The tissue was subsequently stored in 0.1% PFA at 4°C until processing. The tissue was dehydrated, and paraffin embedded. De-waxed sections were stained with anti-UCP1 (ab10983, Abcam), which was detected with VECTASTAIN^®^ ABC kit (VECTOR laboratories).

### 2.6 Indirect calorimetry and activity monitoring

Indirect calorimetry was performed using Home Cage System Phenomaster (TSE Systems). Mice were allowed to acclimate to metabolic chambers for 3-5 days prior to the measurement. Gas exchanges and food intake were recorded every 5 minutes. All metabolic phenotyping data was analyzed by averaging data from 3 days of measurements. The activity of ACBP^-/-^ and K14-ACBP^-/-^ mice were carried out by using implanted telemetric activity monitoring devices. For implantation of telemetric activity monitoring devices, mice were anesthetized with isofluorane and kept on heated pads during the procedure. G2 E-Mitter Telemetry System devices (Starr Life Sciences) were surgically implanted subcutaneously above the shoulder blades and sutured into place. ER4000 Receivers were placed under the cages within the TSE cabinets. The activity of *ma/ma Flg*^*ft/ft*^ mice was monitored by using TSE PhenomasterNG ActiMot3 infrared light beam sensor frames. Physical activity data was integrated into the Phenomaster software.

### 2.7 Statistical analysis

Unless otherwise noted, statistical analysis was performed with GraphPad Prism version 8.4.3. Statistical parameters, including the value of n, are noted in figure legends. Unless otherwise noted, all data are presented as means ± SEM. The statistical significance level was set at p<0.05. For gene expression data an unpaired parametric Student’s t test was used to compare gene expression levels between two genotypes. Repeated measures data was assessed with two-way ANOVA.

## 3. RESULTS

### 3.1 ACBP in keratinocytes is indispensable for normal systemic energy expenditure

We have previously reported that ACBP serves key functions in lipid metabolism in mammals [8; 9] and that ACBP in keratinocytes is required to maintain an intact epidermal barrier and normal fur and to prevent alopecia [10]. To examine how systemic energy homeostasis is affected in ACBP knockout mice, we applied indirect calorimetry to evaluate energy expenditure, respiratory exchange ratio (RER), locomotor activity, and food intake. At room temperature, ACBP^-/-^ mice exhibit significantly increased O_2_ consumption, RER, and food intake compared with wildtype littermates, whereas body weight and locomotor activity remain similar (Figure 1 and Supplementary Fig. S1). This indicates that systemic energy expenditure is increased and that ACBP^-/-^ mice display a substrate preference favoring carbohydrates. Similar to the full body knockout, K14-ACBP^-/-^ mice display increased oxygen consumption and food intake compared with their control littermates (Figure 1), indicating that the increased energy expenditure is linked to the function of ACBP in keratinocytes. Notably however, the RER of K14-ACBP^-/-^ mice remain comparable to that of control mice (Figure 1), showing that the increased RER of ACBP^-/-^ mice is due to the absence of ACBP in other cell types than keratinocytes. Similar to ACBP^-/-^ mice, we did not observe changes in the locomotor activity or body weight of K14-ACBP^-/-^ mice compared with control mice (Figure 1). Taken together, these results show that loss of ACBP expression in keratinocytes leads to an increase in whole body energy expenditure.

**Figure 1.**
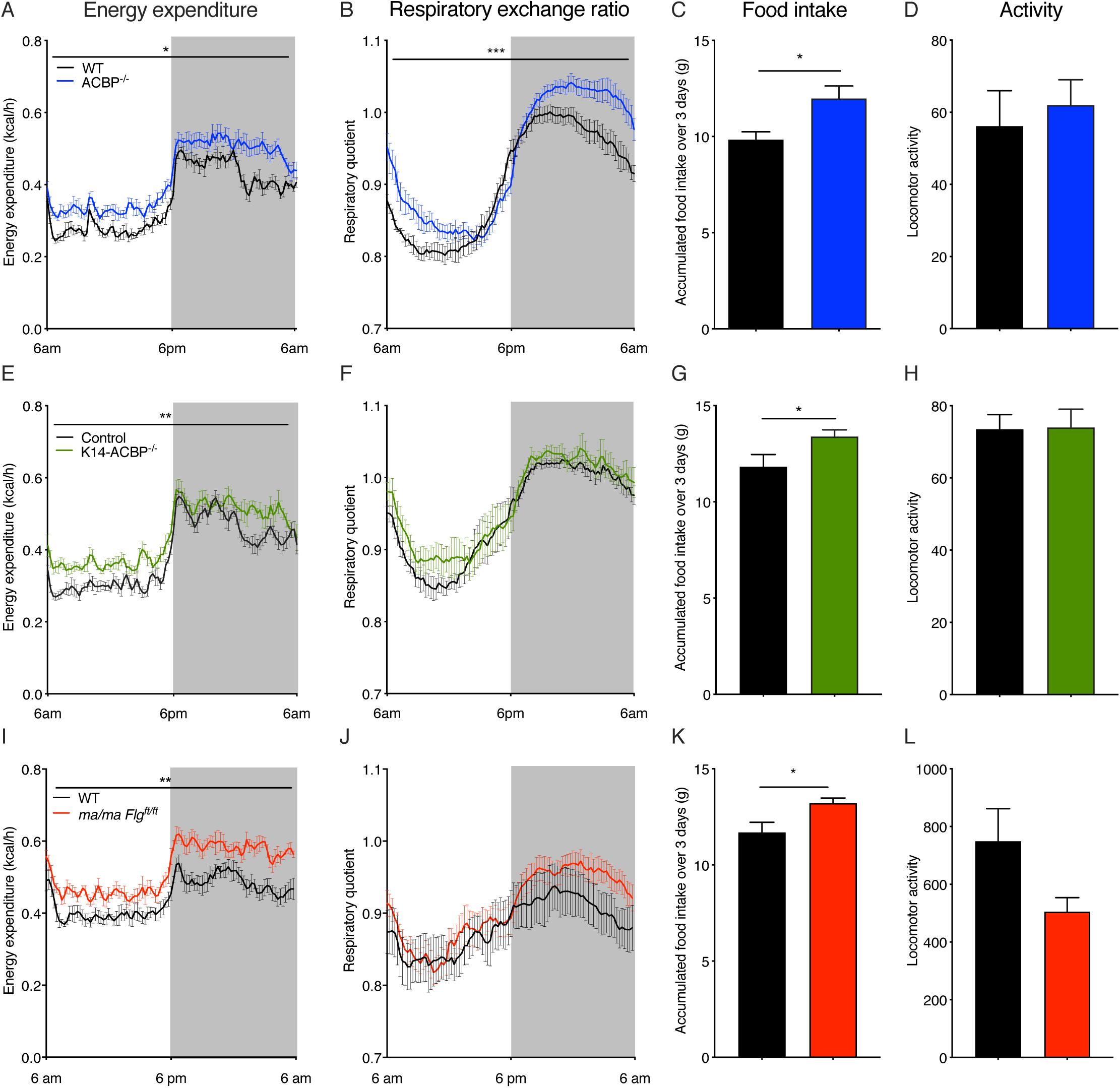
Impaired Epidermal Barrier Leads to Increased Metabolic Rate at room temperature. (A) Energy expenditure at room temperature of wild type (WT) and ACBP^-/-^ mice (ACBP^-/-^) housed in metabolic cages (n = 6 per group, average of 3 days, two-way ANOVA). (B) Daily RQ at room temperature of WT and ACBP^-/-^ mice housed in metabolic cages (n = 6 per group, average of 3 days, two-way ANOVA). (C) Food intake at room temperature of WT and ACBP^-/-^ mice recorded over 3 days in metabolic cages (n=6 per group, Student’s t test). (D) Locomotor activity at room temperature of WT and ACBP^-/-^ mice recorded over 3 days in metabolic cages (n=6 per group, Student’s t test). (E) Energy expenditure at room temperature from control and K14-ACBP^-/-^ mice housed in metabolic cages (n = 6 per group, average of 3 days, two-way ANOVA). (F) Daily RQ at room temperature for control and K14-ACBP^-/-^ mice housed in metabolic cages (n = 6 per group, average of 3 days, two-way ANOVA). (G) Food intake at room temperature of control and K14-ACBP^-/-^ recorded over 3 days in metabolic cages (n=6 per group, Student’s t test). (H) Locomotor activity at room temperature of WT and K14-ACBP^-/-^ mice recorded over 3 days in metabolic cages (n=6 per group, Student’s t test). (I) Energy expenditure at room temperature from WT and *ma/ma Flg*^*ft/ft*^ mice housed in metabolic cages (n = 8 per group, average of 3 days, two-way ANOVA). (J) Daily RQ at room temperature for WT and *ma/ma Flg*^*ft/ft*^ mice housed in metabolic cages (n = 8 per group, average of 3 days, two-way ANOVA). (K) Food intake at room temperature of WT and *ma/ma Flg*^*ft/ft*^ mice recorded over 3 days in metabolic cages (n=8 per group, Student’s t test). (L) Locomotor activity at room temperature of WT and *ma/ma Flg*^*ft/ft*^ mice recorded over 3 days in metabolic cages (n=8 per group, Student’s t test). Data are presented as mean of individuals in each group ± SEM. *p < 0.05; **p < 0.01; ***p < 0.001.

Since transepidermal water loss (TEWL) is significantly increased in both full-body ACBP^-/-^ and K14-ACBP^-/-^ mice [13], we speculated that the increased energy expenditure in these mouse models is caused by the impaired epidermal barrier. To investigate this, we took advantage of another mouse model, the flaky tail mice (*ma/ma Flg*^*ft/ft*^), which carry a frameshift mutation in the filaggrin gene and a mutation in the *Tmem79* gene (*matted, ma*). These mice display increased transepidermal water loss, develop alopecia and are widely used as a model of human atopic dermatitis associated with *FLG* mutations [14; 15]. Interestingly, similar to the ACBP^-/-^ mice, the *ma/ma Flg*^*ft/ft*^ mice exhibit a significantly increased energy expenditure and food intake, while RER is similar to the control mice (Figure 1). Taken together, these findings indicate that defects in the epidermal barrier have major impact on energy metabolism.

### 3.2 A compromised epidermal barrier induces browning of inguinal white adipose tissue

It is well established that prolonged exposure to cold activates thermogenic responses in both humans and rodents, leading to activation of BAT and browning of white adipose tissue (WAT) [16; 17]. To investigate whether the compromised epidermal barrier induces a thermogenic response, we isolated BAT and WAT from ACBP^-/-^, K14-ACBP^-/-^ and *ma/ma Flg*^*ft/ft*^ and control littermates mice housed at room temperature and determined the expression of thermogenic genes. Notably, there is no significant change in UCP1 expression in BAT in any of the mouse models with a compromised epidermal barrier (Supplementary Fig. 2). Interestingly, however, there is a modest but significant induction of thermogenic genes, including uncoupling protein 1 (UCP1), iodothyronine deiodinase 2 (DIO2) and cell death-inducing DNA fragmentation factor-like effector A (CIDEA), in iWAT from ACBP^-/-^ mice, K14-ACBP^-/-^ mice and and *ma/ma Flg*^*ft/ft*^ mice (Figure 2A-C). Consistent with that, the number of multilocular adipocytes in iWAT from ACBP^-/-^ mice, K14-ACBP^-/-^ mice as well as in *ma/ma Flg*^*ft/ft*^ mice is increased compared with iWAT from WT littermates (Figure 2E-F). Furthermore, when the mice were exposed to 4°C for three days, the induction of UCP1 in iWAT was markedly increased in all mouse models with compromised barrier strains compared with their control littermates (Figure 2D-F). These observations indicate that a compromised epidermal barrier leads to an increased thermogenic response in iWAT both at 22°C and at 4°C.

**Figure 2.**
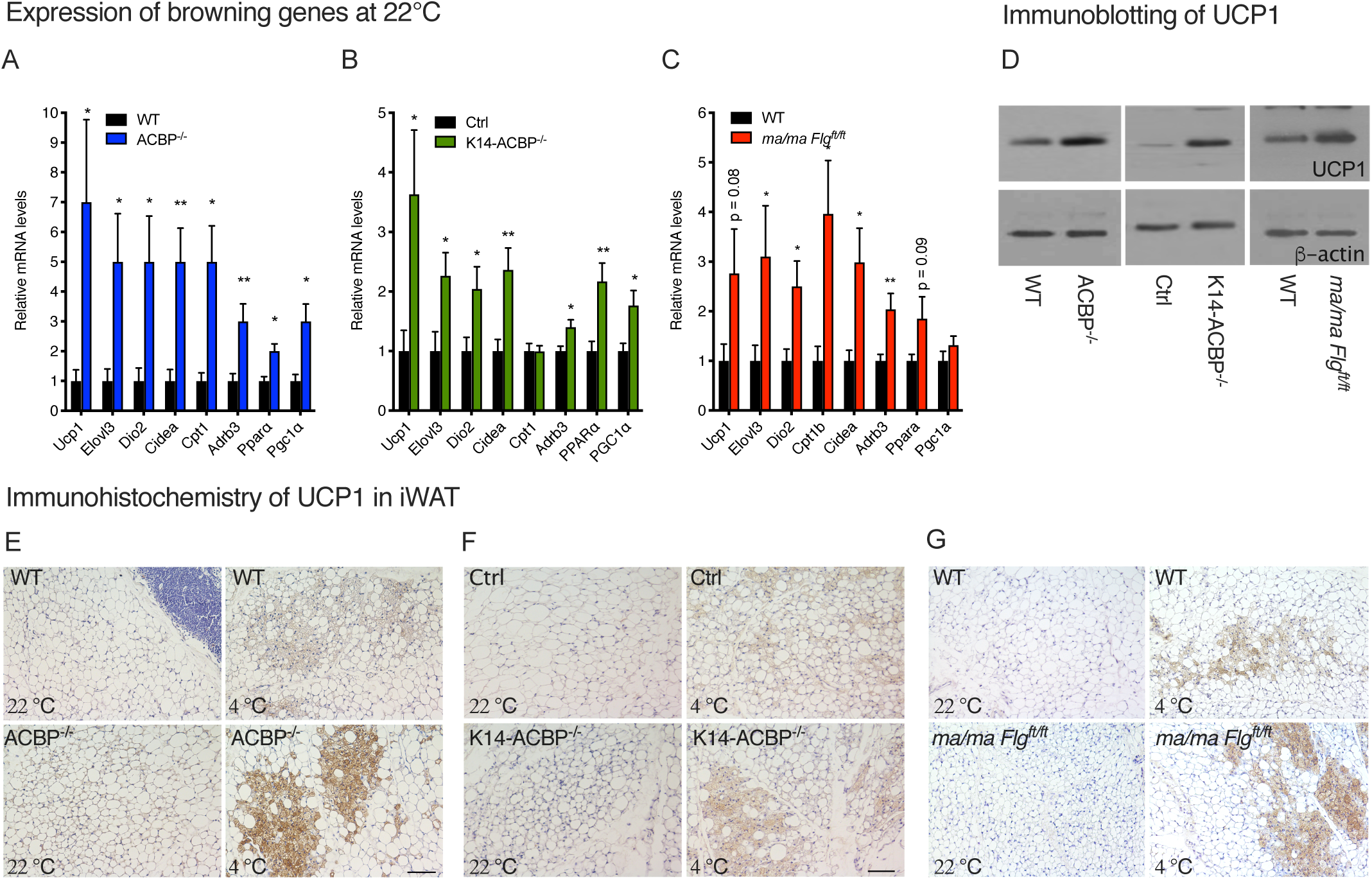
Disruption of ACBP in the Skin Induces Browning of White Adipose Tissue. (A) mRNA levels of *Ucp1, Elovl3, Dio2, Cidea, Cpt1, Adrb3, Ppara* and *Pgc1a* in iWAT of WT and ACBP^-/-^ mice housed at room temperature (n = 12 per group, Student’s t test). Data are presented as mean of individuals in each group ± SEM. *p < 0.05, **p< 0 .01. (B) mRNA levels of *Ucp1, Elovl3, Dio2, Cidea, Cpt1, Adrb3, Ppara* and *Pgc1a* in iWAT of control and K14-ACBP^-/-^ mice housed at room temperature (n = 12 per group, Student’s t test). Data are presented as mean of individuals in each group ± SEM. *p < 0.05, p* < 0.01. (C) mRNA levels of *Ucp1, Elovl3, Dio2, Cidea, Cpt1, Adrb3, Ppara* and *Pgc1a* in iWAT of wildtype and *ma/ma Flg*^*ft/ft*^ mice housed at room temperature (n = 12 per group, Student’s t test). Data are presented as mean of individuals in each group ± SEM. *p < 0.05, p* < 0.01. (D) UCP1 expression determined by Western blotting of iWAT from wildtype, ACBP^-/-^ mice, K14-ACBP^-/-^ and *ma/ma Flg*^*ft/ft*^ mice housed at 4°C for 3 days. Extracts from iWAT from four individual mice were pooled and analyzed by Western blotting and probed for UCP1 (upper panel) and β-actin (lower panel). (E) Representative UCP1 immunostaining of iWAT from control (upper) and ACBP^-/-^ mice (lower) housed at 22°C or 4°C for 3 days, 10x magnification. Scale bars are 100 μm and each picture is representative of 3 individuals. (F) Representative UCP1 immunostaining of iWAT from control (upper) and K14-ACBP^-/-^ mice (lower) housed at 22°C or 4°C for 3 days, 10x magnification. Scale bars are 100 μm and each picture is representative of 3 individuals. (G) Representative UCP1 immunostaining of iWAT from control (upper) and *ma/ma Flg*^*ft/ft*^ mice (lower) housed at 22°C or 4°C for 3 days, 10x magnification. Scale bars are 100 μm and each picture is representative of 3 individuals.

### 3.3 Housing at thermoneutrality rescues systemic energy expenditure and browning of iWAT

The above results indicate that the increased energy expenditure in mice with a compromised epidermal barrier could be due to increased thermogenic activity induced by the increased heat loss. To further investigate this, we housed ACBP^-/-^, K14-ACBP^-/-^ and *ma/ma Flg*^*ft/ft*^ mice and their control littermates in metabolic cages at thermoneutrality. Interestingly, thermoneutrality completely reversed the increased energy expenditure observed in ACBP^-/-^ and K14-ACBP^-/-^ mice at 22°C (Figure 3), whereas energy expenditure in *ma/ma Flg*^*ft/ft*^ mice is not fully rescued at 30°C. Furthermore, the increased food intake of all three epidermal barrier-deficient mouse strains is also reversed when they are housed at 30°C (Figure 3). Similarly, thermoneutrality blunts the induction of browning genes in iWAT of ACBP^-/-^, K14-ACBP^-/-^ and in *ma/ma Flg*^*ft/ft*^ mice compared with their control littermates (Figure 4A-C). These results indicate that the increased thermogenic gene expression in mice with compromised epidermal barrier is caused by heat loss and hence increased cold perception and that increased thermogenic activity leads to an increased energy expenditure and food intake.

**Figure 3.**
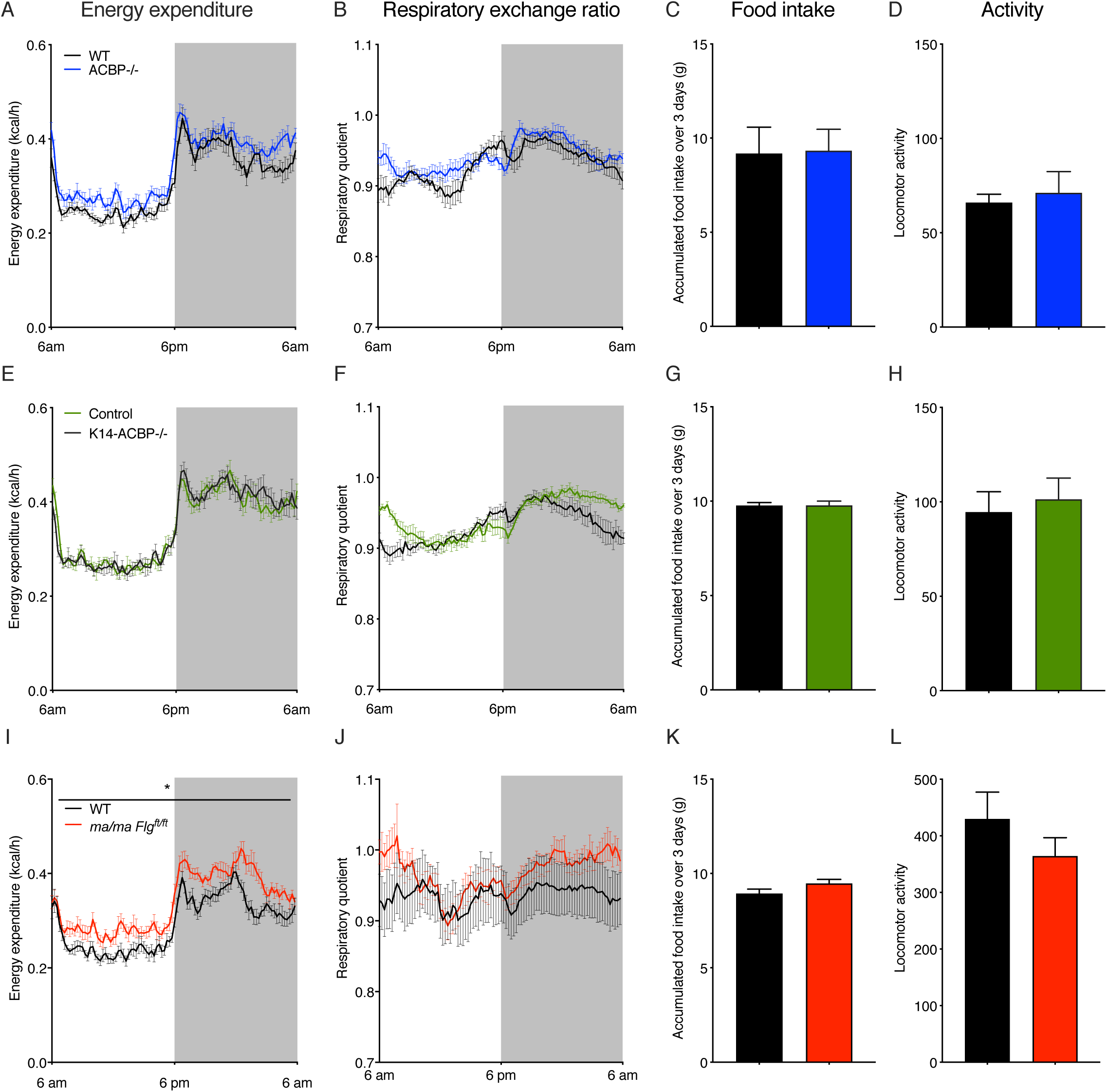
Thermoneutrality Rescues Energy homeostasis of Mice with Compromised Epidermal Barrier. (A) Energy expenditure at 30 °C of WT and ACBP^-/-^ housed in metabolic cages (n = 6 per group, average of 3 days, two-way ANOVA). (B) Daily RQ at 30 °C of WT and ACBP^-/-^ mice housed in metabolic cages (n = 6 per group, average of 3 days, two-way ANOVA). (C) Food intake at 30 °C of WT and ACBP^-/-^ mice recorded over 3 days in metabolic cages (n=6 per group, Student’s t test). (D) Locomotor activity at 30 °C of WT and ACBP^-/-^ mice recorded over 3 days in metabolic cages (n=6 per group, Student’s t test). (E) Energy expenditure at 30 °C from control and K14-ACBP^-/-^ mice housed in metabolic cages (n = 6 per group, average of 3 days, two-way ANOVA). (F) Daily RQ at 30 °C for control and K14-ACBP^-/-^ mice housed in metabolic cages (n = 6 per group, average of 3 days, two-way ANOVA). (G) Food intake at 30 °C of control and K14-ACBP^-/-^ recorded over 3 days in metabolic cages (n=6 per group, Student’s t test). (H) Locomotor activity at 30 °C of WT and K14-ACBP^-/-^ mice recorded over 3 days in metabolic cages (n=6 per group, Student’s t test). (I) Energy expenditure at 30 °C from WT and *ma/ma Flg*^*ft/ft*^ mice housed in metabolic cages (n = 8 per group, average of 3 days, two-way ANOVA). (J) Daily RQ at 30 °C for WT and *ma/ma Flg*^*ft/ft*^ mice housed in metabolic cages (n = 8 per group, average of 3 days, two-way ANOVA). (K) Food intake at 30 °C of WT and *ma/ma Flg*^*ft/ft*^ mice recorded over 3 days in metabolic cages (n=8 per group, Student’s t test). (L) Locomotor activity at 30 °C of WT and *ma/ma Flg*^*ft/ft*^ mice recorded over 3 days in metabolic cages (n=8 per group, Student’s t test). Data are presented as mean of individuals in each group ± SEM. *p < 0.05; **p < 0.01; ***p < 0.001.

**Figure 4.**
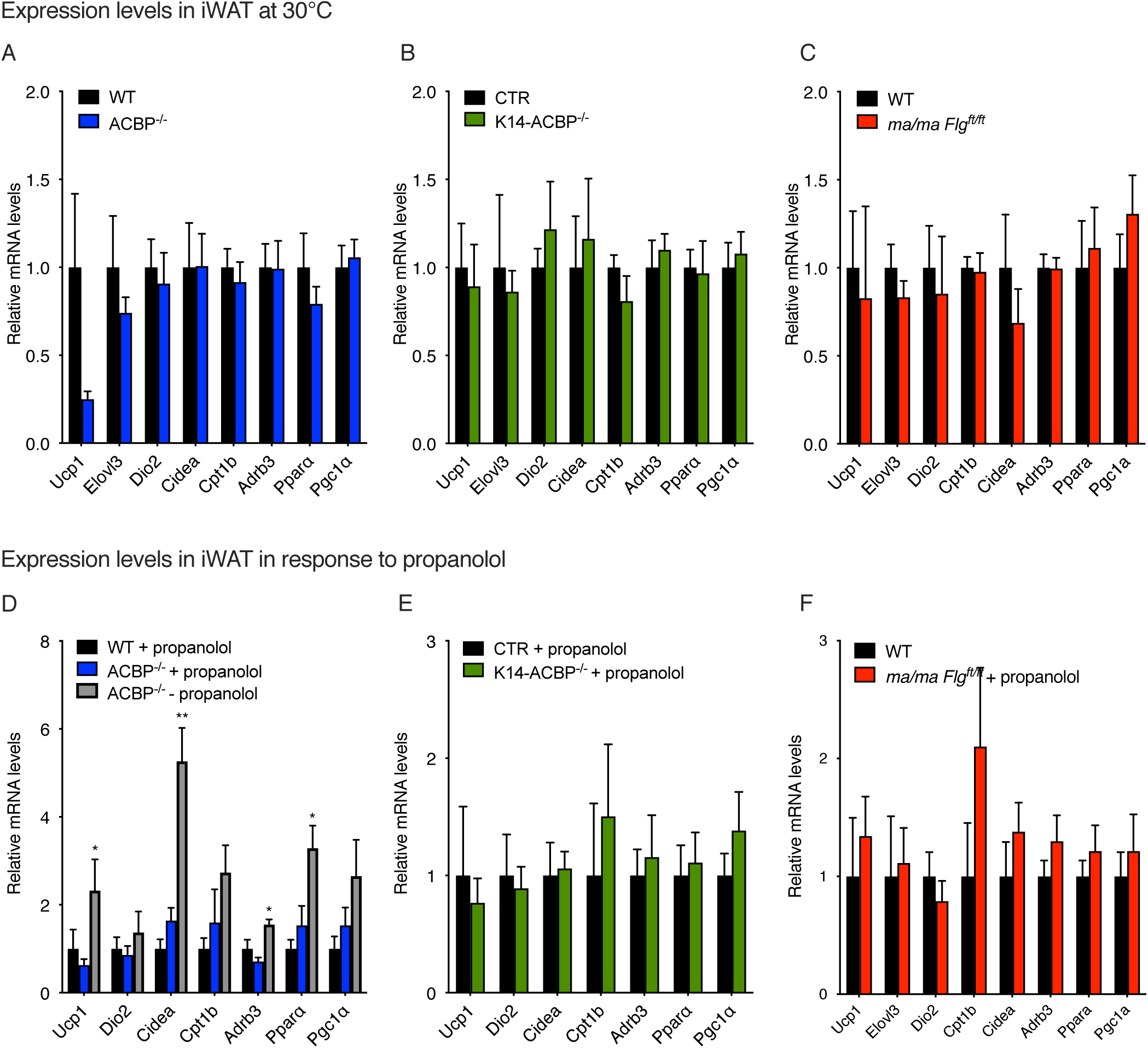
Blocking β-adrenergic Signaling Prevent Browning of Inguinal White Adipose Tissue in ACBP knockout mice. (A) mRNA levels of *Ucp1, Elovl3, Dio2, Cidea, Cpt1, Adrb3, Ppara* and *Pgc1a* in iWAT in WT and ACBP^-/-^ mice housed at thermoneutrality (n = 7-8 per group, Student’s t test). Data are presented as mean of individuals in each group ± SEM. (B) mRNA levels of *Ucp1, Elovl3, Dio2, Cidea, Cpt1, Adrb3, Ppara* and *Pgc1a* in iWAT in WT and K14-ACBP^-/-^ mice housed at thermoneutrality (n = 7-8 per group, Student’s t test). Data are presented as mean of individuals in each group ± SEM. (C) mRNA levels of *Ucp1, Elovl3, Dio2, Cidea, Cpt1, Adrb3, Ppara* and *Pgc1a* in iWAT in WT and *ma/ma Flg*^*ft/ft*^ mice housed at thermoneutrality (n = 7-8 per group, Student’s t test). Data are presented as mean of individuals in each group ± SEM. (D) mRNA levels of *Ucp1, Dio2, Cidea, Cpt1, Adrb3, Ppara* and *Pgc1a* in iWAT of control or propranolol-injected WT and ACBP^-/-^ mice housed at room temperature (n = 5-9 per group, Student’s t test). Data are presented as mean of individuals in each group ± SEM. *p < 0.05, **p< 0 .01 between ACBP^-/-^ mice ± propranolol. (E) mRNA levels of *Ucp1, Dio2, Cidea, Cpt1, Adrb3, Ppara* and *Pgc1a* in iWAT of control or propranolol-injected control and K14-ACBP^-/-^ mice housed at room temperature (n = 5-9 per group, Student’s t test). Data are presented as mean of individuals in each group ± SEM. (F) mRNA levels of *Ucp1, Dio2, Cidea, Cpt1, Adrb3, Ppara* and *Pgc1a* in iWAT of control or propranolol-injected wildtype and *ma/ma Flg*^*ft/ft*^ mice housed at room temperature (n = 5-9 per group, Student’s t test). Data are presented as mean of individuals in each group ± SEM.

The catecholamines adrenaline and noradrenaline control browning and fat cell metabolism mainly through activation of β-adrenoceptors on adipocytes [18]. Since there is little to no sympathetic activity in adipose tissue at thermoneutrality [19], we examined whether the increased browning of iWAT in mice with compromised epidermal barrier at room temperature is dependent on β-adrenoceptor signaling. Hence, we injected mice with propranolol for five days to block β-adrenergic signaling and subsequently examined browning of iWAT and food intake. The results show that propranolol injections blunt the induction of browning genes in ACBP^-/-^, K14-ACBP mice^-/-^ and in *ma/ma Flg*^*ft/ft*^ mice (Figure 4D-F), indicating that the thermogenic response is mediated via β-adrenergic signaling. Moreover, the propranolol injections also reversed the increased food intake of the mouse strains at 22°C (Supplementary Fig. 3). Collectively, our findings show that a compromised barrier function increases thermogenesis and β-adrenergic dependent browning of the iWAT leading to an overall increase in energy expenditure.

### 3.4 Impaired epidermal barrier affords resistance to diet-induced obesity and improves glucose tolerance

Since mice with a compromised epidermal barrier display an elevated energy expenditure, we next sought to investigate whether these mice might also be resistant to diet-induced obesity. Thus, we subjected ACBP^-/-^, K14-ACBP^-/-^, and *ma/ma Flg*^*ft/ft*^ mice and their control littermates to high fat diet (HFD) or standard chow for 12 weeks. On standard chow diet, all mouse models with a compromised epidermal barrier gain weight similar to their control littermates although they eat significantly more than their littermates (Figure 5A,B,D,E,G,H). Interestingly however, all barrier-compromised strains are completely resistant to HFD-induced obesity, although their food intake is comparable to that of control mice (Figure 5).

**Figure 5.**
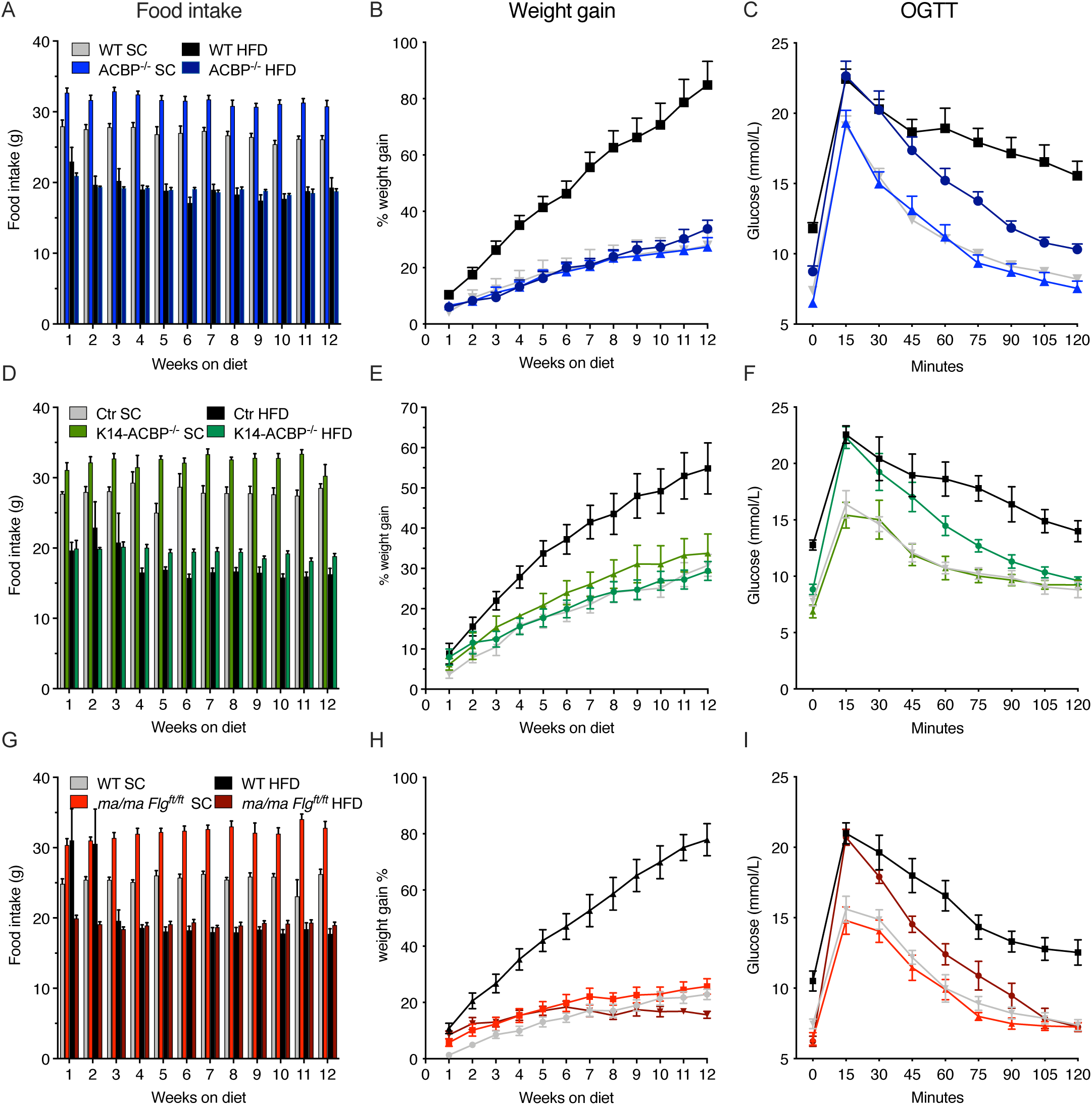
Impaired Epidermal Barrier Protects Mice against Diet-Induced Obesity and Glucose Intolerance. (A) WT and ACBP^-/-^ mice were fed SC or HFD for 12 weeks. Food intake was determined every week for 12 weeks. (B) WT and ACBP^-/-^ mice were fed SC or HFD for 12 weeks. Mice were weighed every week and the percentage weight gain plotted. Two-way ANOVA with multiple comparison was applied. (C) WT and ACBP^-/-^ mice were fed SC or HFD for 12 weeks. Fasting blood glucose was determined prior to administration of 1.5g/kg glucose to each mouse by oral gavage and blood glucose was determined every 15 minutes for a period of 2 hours. (D) Control and K14-ACBP^-/-^ mice were fed SC or HFD for 12 weeks. Food intake was determined was determined every week for 12 weeks. (E) Control and K14-ACBP^-/-^ mice were fed SC or HFD for 12 weeks. Mice were weighed every week and the percentage weight gain plotted. Two-way ANOVA with multiple comparison was applied. (F) Control and K14-ACBP^-/-^ mice were fed SC or HFD for 12 weeks. Fasting blood glucose was determined prior to administration of 1.5g/kg glucose to each mouse by oral gavage and blood glucose was determined every 15 minutes for a period of 2 hours. (G) WT and *ma/ma Flg*^*ft/ft*^ mice were fed SC or HFD for 12 weeks. Food intake was determined was determined every week for 12 weeks. (H) WT and *ma/ma Flg*^*ft/ft*^ mice were fed SC or HFD for 12 weeks. Mice were weighed every week and the percentage weight gain plotted. Two-way ANOVA with multiple comparison was applied. (I) WT and *ma/ma Flg*^*ft/ft*^ mice were fed SC or HFD for 12 weeks. Fasting blood glucose was determined prior to administration of 1.5g/kg glucose to each mouse by oral gavage and blood glucose was determined every 15 minutes for a period of 2 hours. All data are presented as mean of individuals in each group (n=8) ± SEM. *p < 0.05; **p < 0.01; ***p < 0.001.

These observations prompted us to examine whether mice with a compromised epidermal barrier are also protected from developing insulin resistance on HFD. We therefore subjected ACBP^-/-^, K14-ACBP^-/-^ and *ma/ma Flg*^*ft/ft*^ mice fed either a HFD or chow for 12 weeks to an oral glucose tolerance test (OGTT) (Figure 5C,F,I). On chow diet, fasting glucose levels and glucose clearance are similar between ACBP^-/-^, K14-ACBP^-/-^, and *ma/ma Flg*^*ft/ft*^ mice and their respective control littermates. However, on HFD all mouse models with a compromised epidermal barrier are protected from the HFD-induced increase in fasting glucose concentrations and display an increased glucose clearance relative to their control littermates (Figure 5C,F,I), indicating that these mice are partially protected from developing insulin resistance on HFD. Notably however, despite similar body weight, glucose clearance is lower in HFD compared with chow fed mutant mice, indicating that the HFD *per se*, leads to a compromised glucose tolerance independent of obesity. This does not appear to be caused by insulin resistance, since ACBP^-/-^ and K14-ACBP^-/-^ mice on HFD display plasma insulin levels comparable to that of mice on chow, whereas the *ma/ma Flg*^*ft/ft*^ mice on HFD have significantly decreased plasma insulin levels compared with control mice (Figure 6).

**Figure 6.**
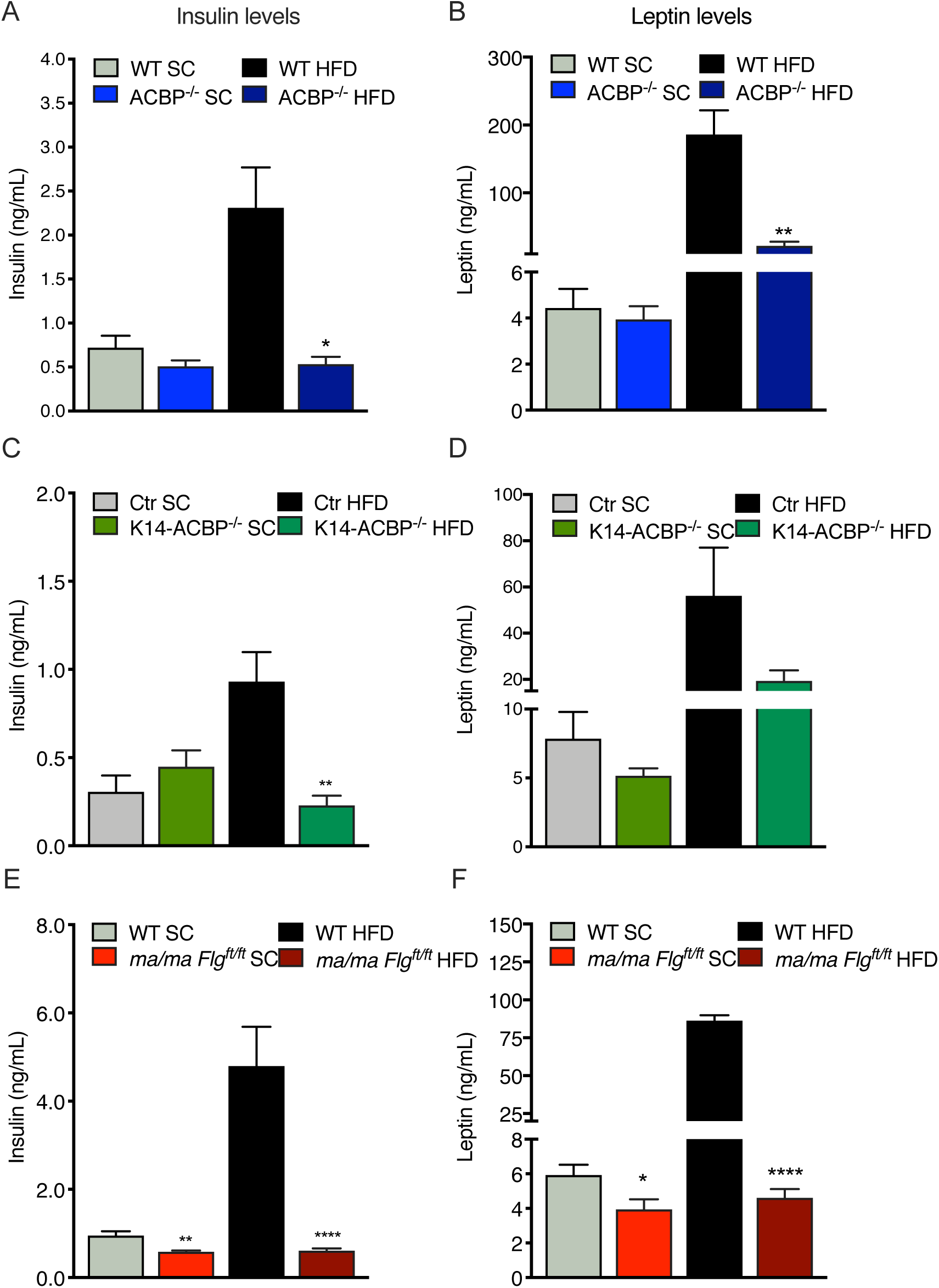
Impaired Epidermal Barrier Protects Mice against Diet-Induced Hyperinsulinemia. (A) Plasma insulin levels in WT and ACBP^-/-^ mice fed either SC or HFD for 12 weeks. (B) Plasma leptin levels in WT and ACBP^-/-^ mice fed either SC or HFD for 12 weeks. (C) Plasma insulin levels in control and K14-ACBP^-/-^ mice fed either SC or HFD for 12 weeks. (D) Plasma leptin levels in control and K14-ACBP^-/-^ mice fed either SC or HFD for 12 weeks. (E) Plasma insulin levels in WT and *ma/ma Flg*^*ft/ft*^ mice fed either SC or HFD for 12 weeks. (F) Plasma leptin levels in WT and *ma/ma Flg*^*ft/ft*^ mice fed either SC or HFD for 12 weeks. All data are presented as mean of individuals in each group (n=8) ± SEM. *p < 0.05; **p < 0.01; ***p < 0.001.

Plasma leptin levels are known to show a strong positive correlation with the amount of total body fat [20; 21]. We found that ACBP^-/-^ and K14-ACBP^-/-^ mice have plasma leptin concentrations comparable to that of the WT mice, whereas *ma/ma Flg*^*ft/ft*^ mice have significantly decreased leptin levels. Consistent with the absence of HFD-induced obesity in mice with compromised epidermal barrier, plasma leptin levels in these mice are lower than in the control littermates when fed HFD (Figure 6B,D,F). Collectively, our data show that despite a similar food intake, mice with a compromised epidermal barrier are resistant to HFD-induced obesity and maintain normal fasting glucose and insulin levels on HFD.

## 4. DISCUSSION

In the present study we show that mice with impairment in epidermal barrier function in the ACBP^-/-^, K14-ACBP^-/-^ and *ma/ma Flg*^*ft/ft*^ models display increased energy expenditure that appear to be driven by increased thermogenesis. These mice are also completely protected from HFD-induced obesity, and partially protected from the HFD-induced decrease in glucose clearance.

We have previously shown that ablation of ACBP in the skin diminishes very long-chain fatty acid levels in the stratum corneum, which are required for epidermal barrier function [10]. As a consequence of this, the sebaceous as well as the Harderian glands become hyperplastic and produce abnormal high levels of lipids, which are believed to be secreted to provide further insulation ([13] and our unpublished results). The present findings show that ACBP^-/-^ and K14-ACBP^-/-^ mice in addition upregulate their energy expenditure to defend their body temperature when housed at room temperature. The increased energy expenditure of ACBP^-/-^ and K14-ACBP^-/-^ mice is so profound that the mice are resistant to diet-induced obesity and the diabetogenic effects of a high-fat diet. This is contrary the effect of ablation of ACBP in astrocytes in the arcuate nucleus, which has been shown to promote diet-induced hyperphagia and obesity in mice [22]. Thus, systemic effects resulting from loss of ACBP in keratinocytes appear to override the effects of ACBP in astrocytes in control of energy balance and its anorectic effects.

Interestingly, we found that ACBP^-/-^ mice have an increased RER compared to WT mice, whereas RER of K14-ACBP^-/-^ and *ma/ma Flg*^*ft/ft*^ mice is unchanged, indicating that the rise in RER is not caused by the increased energy metabolism. Instead the rise in RER indicates that lack of ACBP in other cell types than keratinocytes leads to an increase in the ratio of carbohydrate to fatty acid oxidation. This finding is in line with the observation that ACBP is required to sustain fatty acid oxidation in glioma cells [23] and in human lung cancer cells [24] and is able to mediate transport of long-chain acyl-CoA esters to mitochondria *in vitro* [25].

Other mouse models with impaired synthesis of epidermal lipids support the notion that epidermal lipid metabolism plays a major role in epidermal barrier function. For instance, loss of elongation of very long chain fatty acid (ELOVL3), diacylglycerol O-acyltransferase 1 (DGAT1), stearoyl-CoA desaturase (SCD1) or alkaline ceramide (ACER1) also impair synthesis of epidermal barrier lipids in mice, enhance evaporative cooling, increase whole-body energy expenditure and render the mice resistant to diet-induced obesity [4; 5; 7; 26; 27]. To substantiate the role of the epidermal barrier in systemic energy metabolism, we also show that mice with functional loss of filaggrin and Tmem79, which are not involved in lipid metabolism *per se* [28], share several of the phenotypic characteristics of the ACBP^-/-^ mice including increased TEWL, tousled fur, alopecia, and enlarged Harderian glands ([14; 15] and our unpublished results) and exhibit elevated energy expenditure. Therefore, besides epidermal lipid metabolism, loss of structural entities like filaggrin, are also required to maintain systemic energy metabolism.

This notion is in keeping with recent observations showing that a compromised epidermal barrier augments evaporative cooling at the body surface, which further enhances energy expenditure that accounts for a significant proportion of total systemic energy metabolism in mice [29]. Moreover, shaving of wild type mice increases the metabolic rate at room temperatures (approximately 22°C) by approximately 50% due to increased heat loss [30], which nearly corresponds to transferring an unshaved mouse from 20°C to 10°C [31]. Furthermore, Nedergaard and Cannon stated that at 20°C, the extra metabolism required to counteract the extra heat loss due to shaving is nearly 3 times as high than that of an unshaved mouse. Additionally, genetically nude Balb/c mice have a metabolic rate, which is almost 2 fold higher than that of normal Balb/c mice [30]. Thus, impaired insulation caused by changes in the quality of the fur and hair, resembles exposure to colder ambient temperatures.

Cold exposure leads to a profound increase in whole body energy expenditure of small rodents, and that activation of non-shivering thermogenesis in brown adipose tissue (BAT) plays a key role in in defending body temperature during prolonged cold exposure [32]. In this study, expression of UCP1 was neither increased in BAT of full-body ACBP^-/-^, skin-specific K14-ACBP^-/-^ mice, nor in *ma/ma Flg*^*ft/ft*^. However, we consider it likely that increased thermogenic activity in BAT, induced by non-transcriptional mechanisms, plays a major role in the increased energy expenditure in the mice with the compromised epidermal barrier. Analysis of iWAT demonstrated modest browning at room temperature and a more pronounced browning at 4°C in all three mouse models with compromised epidermal barrier compared with their littermate controls. Further analyses indicated that β-adrenergic signals are likely to play a major role in browning of iWAT, since blocking of β-adrenergic signaling by propranolol injections and housing at thermoneutrality completely alleviated browning of iWAT and normalized food intake. In line with this observation, it was recently demonstrated that a β3-adrenergic receptor agonist not only improves oral glucose tolerance and insulin sensitivity but also stimulates browning of subcutaneous WAT in obese humans [33].

To what extent the epidermal barrier contributes to regulation of systemic energy metabolism in humans is not fully resolved. Not only is the structure and morphology of the skin different between mice and humans, the surface to volume ratio is much greater for small rodents than for humans and the adipose tissue is far less thermogenic in humans compared with rodents. Nevertheless, it is interesting to note that humans suffering from burn injuries or children suffering from ichthyosis exhibit not only increased transepidermal barrier disruption but also elevated resting energy expenditure [34-36]. We have shown that loss of ACBP in keratinocytes and the ensuing impairment of epidermal barrier function leads to a profound increase in energy metabolism, browning of white adipose tissue and protection against HFD-induced obesity, phenotypes that are recapitulated in *ma/ma Flg*^*ft/ft*^ mice. Our results indicate that an increase in epidermal barrier permeability, can enhance systemic energy expenditure and hence protect against an increased accumulation of calories and eventually metabolic diseases.

## Author contributions

D.N., A.-B. M., Z.G.-H., S.M., and N.J.F. conceived and designed the project; V.K, D.N, A.-B.M., M.R.W. J.V., P.M.M., J.R.B, R.P., G.C, I.S., T.M, and I.S. performed the experiments; V.K., D.N., A.-B.M., Z.G.-H., M.R.W., J.V., P.M.M., R.P., T.M, G.C, I.S., S.M., and N.J.F. analyzed and interpreted data; D.N. and N.J.F. wrote the original draft; D.N., A.-B.M., S.M., and N.J.F. wrote the final version.

## Acknowledgements

The skillful technical assistance of Ida Nørgaard Jensen is gratefully acknowledged.

## Figure legends

**Supplementary Figure S1. Body weights of 7-8 weeks old mice**

(A) Body weights of in WT and ACBP^-/-^ mice housed at room temperature (n = 6-8 per group). Data are presented as mean of individuals in each group ± SEM.

(B) Body weights of in control and K14-ACBP^-/-^ mice housed at room temperature (n = 6-8 per group). Data are presented as mean of individuals in each group ± SEM.

(C) Body weights of in WT and *ma/ma Flg*^*ft/ft*^ mice housed at room temperature (n = 6-8 per group). Data are presented as mean of individuals in each group ± SEM.

**Supplementary Figure S2. UCP1 expression in brown adipose tissue in epidermal barrier compromised mice**

(D) mRNA levels of *Ucp1* in BAT in WT and ACBP^-/-^ mice housed at room temperature (n = 7-8 per group, Student’s t test). Data are presented as mean of individuals in each group ± SEM.

(E) mRNA levels of *Ucp1* in BAT in control and K14-ACBP^-/-^ mice housed at room temperature (n = 7-8 per group, Student’s t test). Data are presented as mean of individuals in each group ± SEM.

(F) mRNA levels of *Ucp1* in BAT in WT and *ma/ma Flg*^*ft/ft*^ mice housed at room temperature (n = 7-8 per group, Student’s t test). Data are presented as mean of individuals in each group ± SEM.

**Supplementary Figure S3. Blocking** β**-adrenergic signaling at 22**°C **normalizes food intake in epidermal barrier compromised mice**

(A) Food intake at 22°C of wild type and ACBP^-/-^ mice recorded over 3 days before and after injection of propanolol (n=7 per group, Student’s t test). Data are presented as mean of individuals ± SEM.

(B) Food intake at 22°C of control and K14-ACBP^-/-^ recorded over 3 days before and after injection of propanolol (n=7 per group, Student’s t test). Data are presented as mean of individuals ± SEM.

(C) Food intake at 22°C of control and *ma/ma Flg*^*ft/ft*^ mice recorded over 3 days before and after injection of propanolol (n=7 per group, Student’s t test). Data are presented as mean of individuals ± SEM. *p < 0.05; **p < 0.01; ***p < 0.001.

**Table S1.**
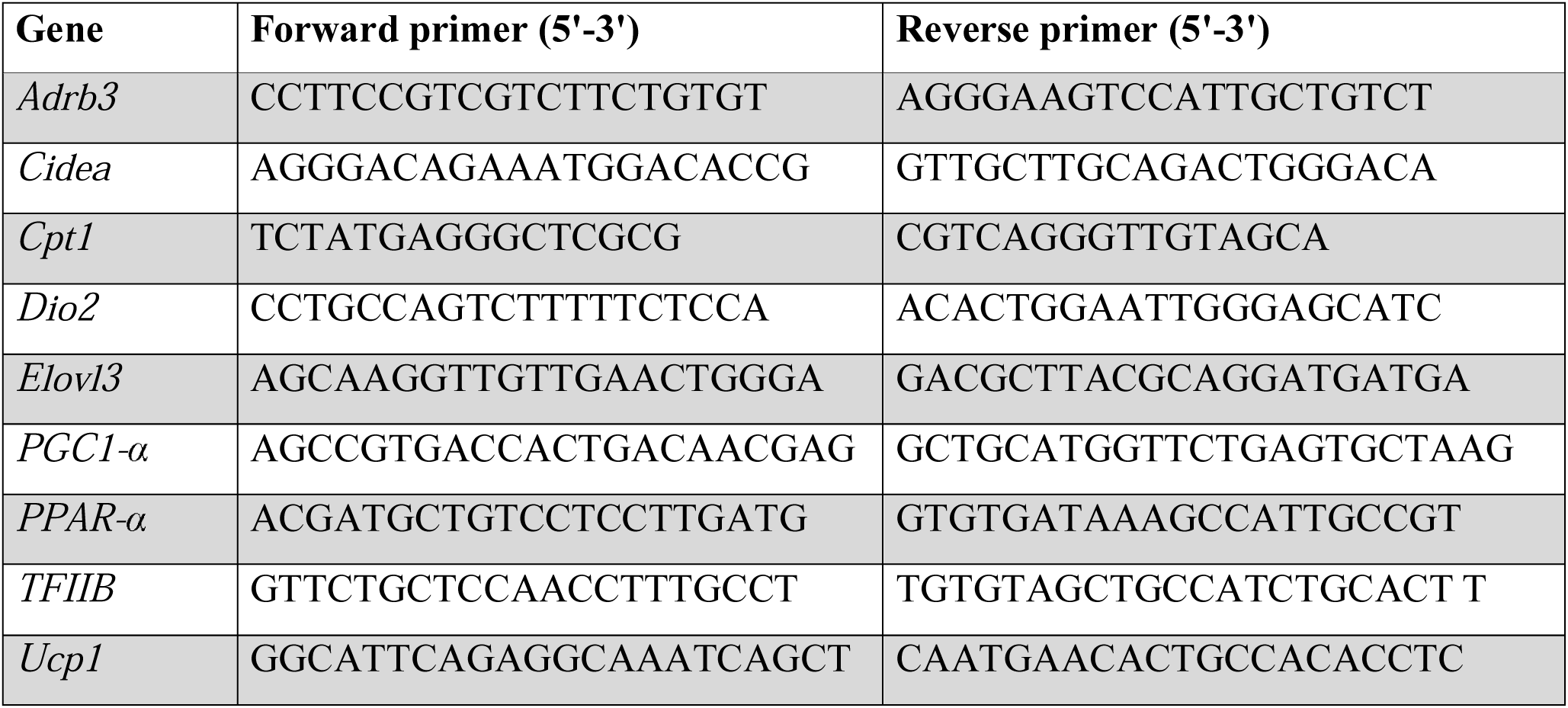
Sequences of qPCR primers used.

