## Supplementary figures and images for "The Epidermal Barrier is Indispensable for Systemic Energy Homeostasis"

### Supplemental figure 1

Figure S1

Mouse body weights

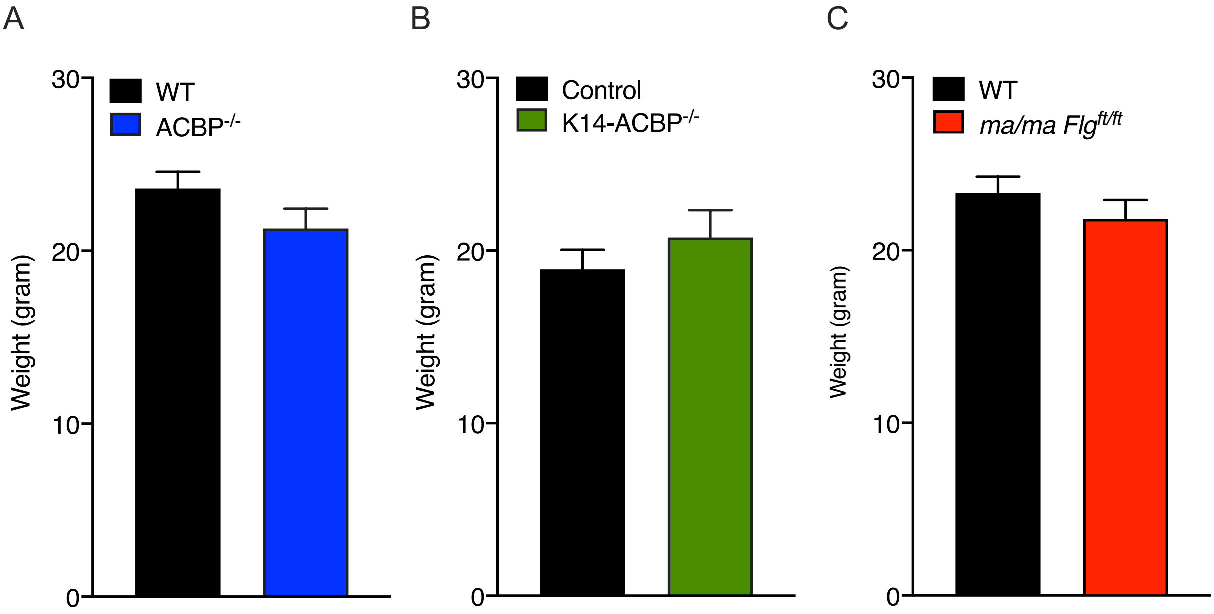

### Supplemental figure 2

Figure S2

Expression of UCP1 in BAT

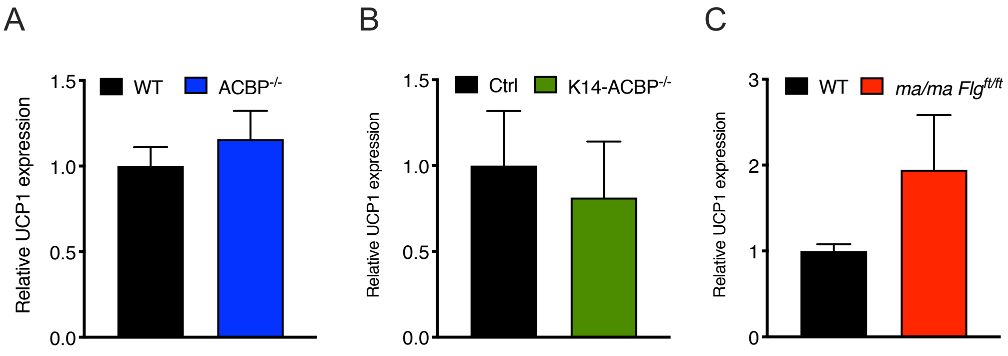

### Supplemental figure 3

Figure S3

Food intake before and after propanolol injections

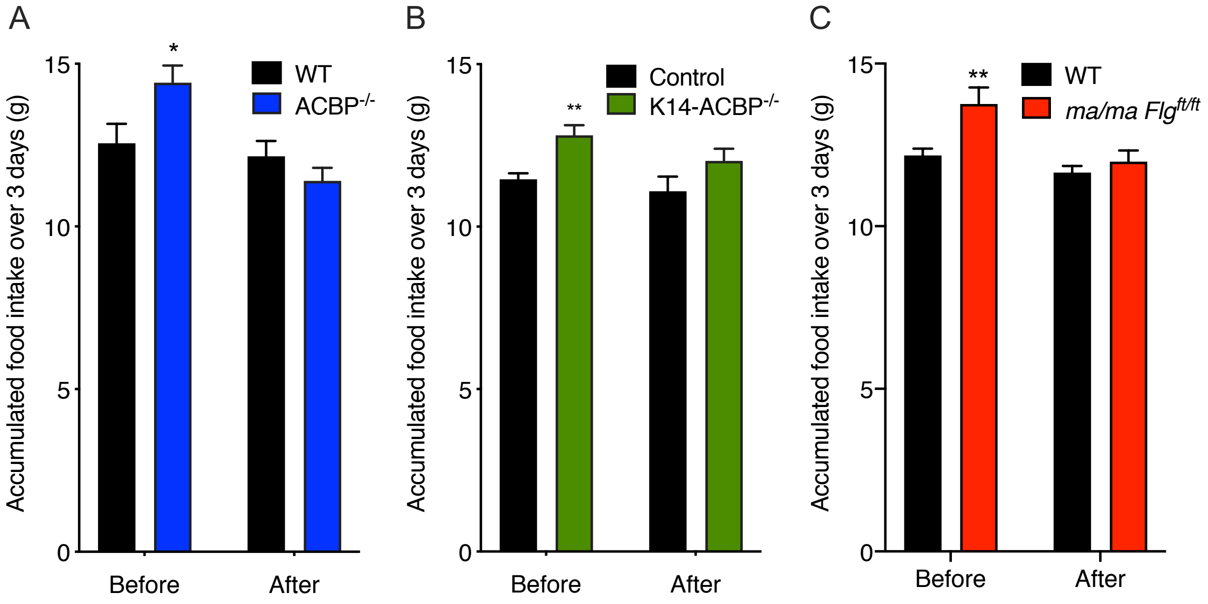
